# Molecular patterning during the development of *Phoronopsis harmeri* reveals similarities to rhynchonelliform brachiopods

**DOI:** 10.1101/782839

**Authors:** Carmen Andrikou, Yale J. Passamaneck, Chris J. Lowe, Mark Q. Martindale, Andreas Hejnol

## Abstract

**Background:** Answering the question how conserved patterning systems are across evolutionary lineages requires a broad taxon sampling. Phoronid development has previously been studied using fate mapping and morphogenesis, yet molecular descriptions are missing. Here we report the expression patterns of the evolutionarily conserved anterior (*otx, gsc, six3/6, nk2.1*), posterior (*cdx, bra*) and endomesodermal (*foxA, gata4/5/6, twist*) markers in the phoronid *Phoronopsis harmeri*.

**Results:** The transcription factors *foxA, gata4/5/6* and *cdx* show conserved expression in patterning the development and regionalization of the phoronid embryonic gut, with *foxA* expressed in the presumptive foregut, *gata4/5/6* demarcating the midgut and *cdx* confined to the hindgut. Surprisingly, *brachyury*, an evolutionary conserved transcription factor often associated with gastrulation movements and patterning of the mouth and hindgut, seems to be unrelated with gastrulation and mouth patterning in phoronids. Furthermore, *six3/6*, a well-conserved anterior marker, shows a remarkably dynamic expression, demarcating not only the apical organ and the oral ectoderm, but also clusters of cells of the developing midgut and the anterior mesoderm, similar to what has been reported for brachiopods, bryozoans and some deuterostome Bilateria.

**Conclusions:** Our comparison of gene expression patterns with other studied Bilateria reveals that the timing of axis determination and cell fate distribution of the phoronid shows highest similarities to rhynchonelliform brachiopods. Despite these similarities, the phoronid *P. harmeri* shows also particularities in its development, which hint to divergences in the arrangement of gene regulatory networks responsible for germ layer formation and axis specification.

## Background

Phoronids are small, filter-feeding, sessile marine worms that are placed by molecular phylogenetic analyses together with the Brachiopoda and Ectoprocta in a clade called Lophophorata [1–5], Phoronida is subdivided in two main taxa, *Phoronis* Wright 1856 and *Phoronopsis* Gilchrist 1907, to which one species, *Phoronis ovalis*, forms the sister species (Figure 1A) [6], Most phoronids are characterized by a planktotrophic actinotroch larva (Figure 1B), which undergoes a rapid, catastrophic metamorphosis in order to give rise to the adult body plan [7, 8]. The development of phoronids has been described by a number of authors and, except for differences in the cleavage pattern and the mode of coelom formation, appears to be similar between species [9–22], Cleavage is holoblastic, and the first two divisions are meridional along the animal-vegetal main axis [9–12, 16–20, 22, 23], At the eight-cell stage, the embryo is composed of an animal and a vegetal tier of four cells, but the blastomeres vary in their arrangement between embryos [9–12, 16–20, 22, 23], The variability is also seen in the next division rounds and led some authors to describe the phoronid cleavage pattern as radial or biradial [10, 12, 14–17, 22, 24], spiral [9, 18, 19, 25, 26], or even a transition between a radial and spiral pattern [27, 28], By the 64-cell stage, the embryo develops into a ciliated blastula [12, 16, 18–20, 22–24, 28], Blastulae can be thick-walled with a small blastocoel (e.g. *Phoronis psammophila*) [24] or thin-walled with an extensive blastocoel (e.g. *Phoronopsis harmeri*) [19, 22], Gastrulation begins with the flattening of the vegetal pole of the embryo and the subsequent invagination of the archenteron, that forms a centrally located blastopore [12, 14, 16, 19, 20, 22, 24, 28], In phoronids, the animal-vegetal axis of the early embryo does not correspond to the anterior-posterior axis of the larva [12, 14, 16, 19, 20, 22, 24, 28], During gastrulation, both the animal pole and the blastopore shift towards the future anterior end of the larva, whilst the embryo and the developing archenteron elongate in an anterior to posterior direction, establishing the future plane of the bilateral symmetry of the larva [12, 14, 16, 19, 20, 22, 24, 28]. An anterior ectodermal thickening leads to the formation of the apical organ [12, 14, 16, 19, 20, 22, 24, 28], At the end of gastrulation, the blastopore is reduced to a round-shaped anterior remnant that will form the mouth, while the anus will open independently at the posterior end of the larva [12, 14, 16, 19, 20, 22, 24, 28]. Mesoderm originates in two waves: from an anterior domain of the invaginating archenteron at early gastrula stage and from a posterior ventrolateral domain of the elongated archenteron at larva stage [9–14, 19–21, 25], The anterior mesoderm will form the cavity that fills the pre-oral lobe and the muscles of the pre-oral lobe, and the posterior mesoderm will form the trunk coelom (metacoel) [9–14, 19–21, 25], The formation of the pre-oral lobe cavity shows variation between species. Mesodermal cells can proliferate and form the lining of a coelom (protocoel) (e.g. *Phoronis ijimai, Phoronis psammophila* and *Phoronopsis harmeri*) [9, 15, 21, 24, 28–30], or can form a cell mass that is surrounded by extracellular matrix (e.g. *Phoronis muelleri*) [31].

**Figure 1.**
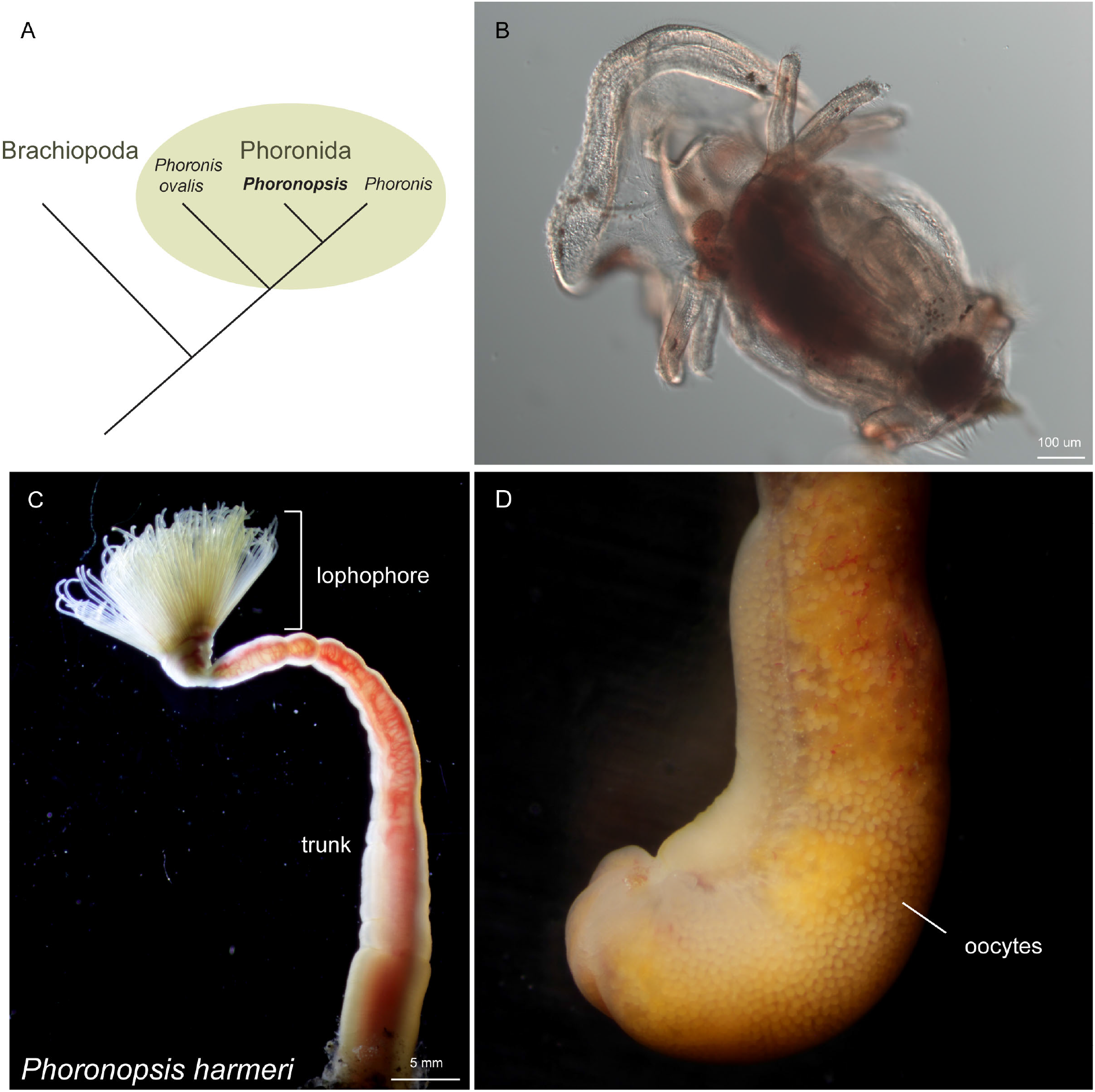
Gross morphology and phylogenetic position of *Phoronopsis harmeri*. **(A)** Phylogenetic position of phoronids [6]. **(B)** Characteristic actinotroch larva with 12 tentacles. **(C)** Anterior region of *Phoronopsis harmeri* with a terminal anterior lophophore, used for collection of food particles and respiration, and a posterior trunk. **(D)** Ampulla of a mature female animal with visible oocytes. Anterior is to the top.

Blastomere ablation experiments on *Phoronis ijimai* and *Phoronopsis harmeri* have demonstrated the large regulative potential of phoronids, since blastomeres isolated at the 2-cell stage are able to produce complete, but diminutive embryos [22]. Moreover, fate-mapping experiments in *Phoronis vancouverensis* have shown that the early animal tier of the eight-cell embryo forms only ectoderm, while the early vegetal tier forms ectoderm, endoderm and mesoderm [12]. A later study on the same species suggested that muscles and neurons originate from portions of endoderm and ectoderm, and that the intestine forms by ingression of the posterior ectoderm [13]. However, molecular studies on embryonic development of phoronids are still lacking and are important to understand the precise timing of germ layer segregation and cell specification.

In this study we investigated the embryonic gene expression of the phoronid *Phoronopsis harmeri* Pixell, 1912. *P. harmeri* occurs in very large numbers in coastal intertidal mudflats of the North Pacific. The body of the adult animal is subdivided into two main compartments, an anterior lophophore and a posterior trunk (Figure 1C) with a terminal ampulla (Figure 1D) [32]. Fertilization takes place internally in the coelomic fluid of the female trunk (Figure 1D) and each gravid adult can release hundreds of eggs. The cleavage pattern of *P. harmeri* is a debated subject; some authors consider it radial [20, 22, 28] and others spiral (referred as *P. viridis* in [19]).

To identify the appearance and segregation of the primary embryonic fates along the anterior-posterior axis in *P. harmeri*, we first identified orthologs of evolutionary conserved genes mainly associated with anterior (*otx, gsc, six3/6, nk2.1*) [33–37], posterior (*cdx, bra*) [38, 39] and endomesodermal identities (*foxA, gata4/5/6, twist*) [40–45], and revealed their spatial expression during embryonic development by Whole Mount In Situ Hybridization (WMISH). By comparing our findings with brachiopods, the proposed sister group of phoronids, we show that *P. harmeri* shares more similarities in molecular patterning to rhynchonelliform (e.g. *Terebaratalia transversa*), than craniiform (e.g. *Novocrania anomala*) brachiopods. Our analysis provides a description of marker gene expression during phoronid development and highlights clear cases of conservation but also novelties of embryonic gene expression.

## Results

### Embryological description of the development of *P. harmeri*

To better understand the spatial and temporal expression of the candidate developmental genes, we first analyzed the developmental stages of *P. harmeri* using differential interference contrast (DIC) and confocal laser scanning microscopy.

After fertilization, two polar bodies are formed; the first polar body is formed soon after the release of the eggs into the seawater, and the next about 30 minutes later, both of which remain associated with the embryo due to the presence of a thick vitelline membrane. The first division is meridional and occurs approximately 2 hours after the egg contacts the seawater (Figure 2A). The second cleavage is also meridional but perpendicular to the preceding division and takes place one hour after the completion of the previous division (Figure 2B). The third division starts around 30-60 minutes later in the equatorial plane, with the blastomeres of the animal quartets oriented directly above the vegetal ones (Figure 2C).

**Figure 2.**
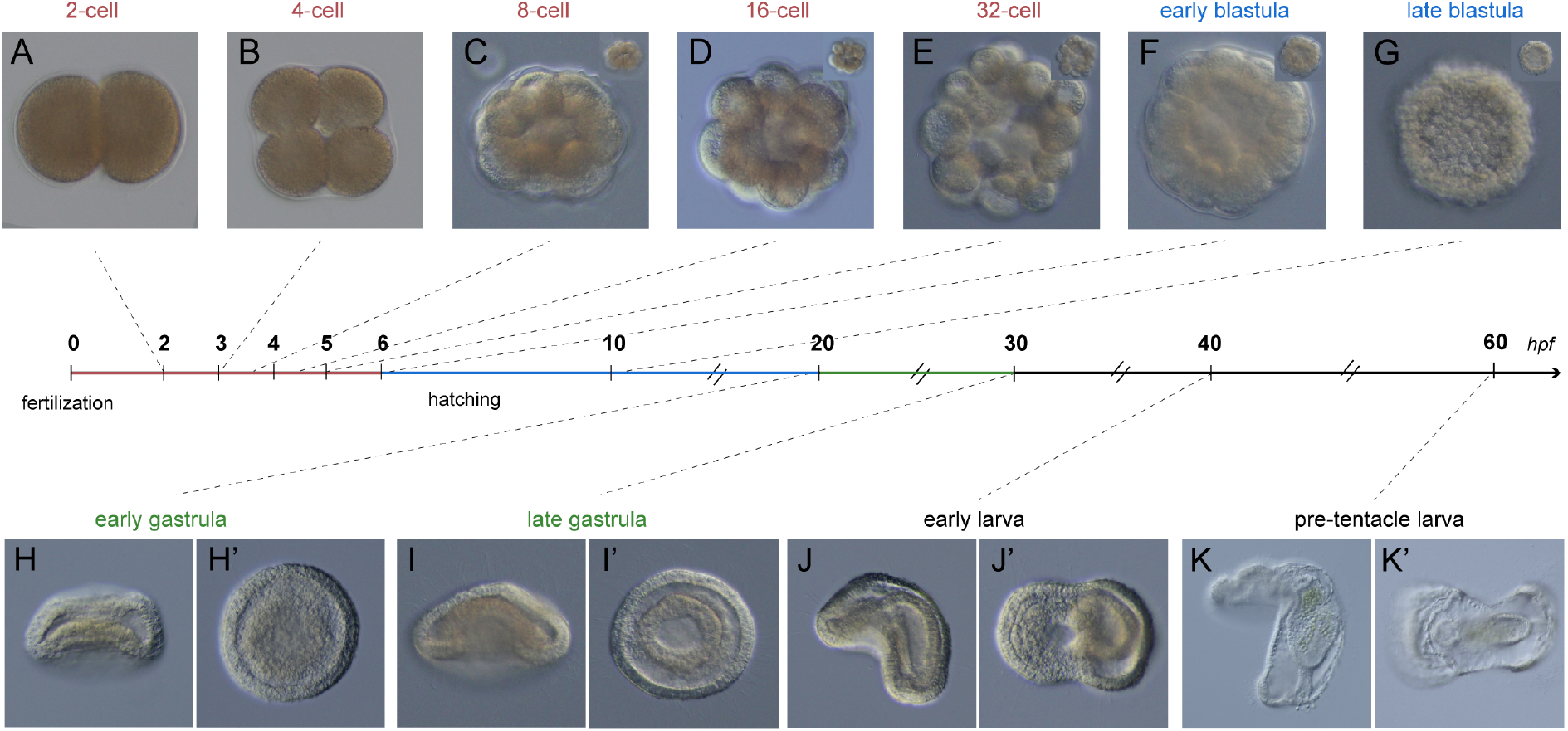
The embryonic development of *P. harmeri*. Nomarski images of living embryos of *P. harmeri* at representative stages of development. The egg undergoes its first radial holoblastic cleavage 2hpf and forms a hatching blastula around 6-10hpf. Gastrulation starts at 20hpf at the vegetal pole of the embryo and results in the flattening of the vegetal surface. At late gastrula stage (30hpf) the apical organ shifts anteriorly, the archenteron elongates posteriorly and the anterior-posterior axis becomes oblique. At early larva stage (40hpf) the embryo begins to elongate along the anterior-posterior axis and the blastopore becomes the mouth of the future larva. A thick tissue is formed at the dorsal ectoderm and around the mouth that will form the future pre-oral lobe. The bilateral symmetry is evident. The pre-tentacle actinotroch larva is formed around 60hpf, with a prominent pre-oral lobe, a fully compartmentalized, functional gut and evident tentacle bulbs. Panels H, I, J and K depict embryos in lateral view and panels H’, I’, J’ and K’ show embryos in vegetal view. Insets show different focal planes of the embryos. In all panels, anterior is to the left.

This third cleavage and the next two divisions result in the formation of different blastomere arrangements of spiral-like appearance (Figure 2C-E). A ciliated blastula with cone-shaped cells is formed at approx. 6-8 hours post fertilization (hpf) (Figure 2F). Within the next couple of hours, the blastula hatches and starts to swim. Around lOhpf, a large blastocoel is evident (Figure 2G and 3A). The onset of gastrulation occurs at approximately 20hpf (early gastrula stage) with a flattening of the vegetal pole and the formation of a shallow indentation (Figure 2H and 3B). At 30hpf (late gastrula stage), cells ingress in the blastocoel and the archenteron epithelium thickens and elongates, due to the axial elongation of the embryo. The ciliated apical organ shifts approximately 90 degrees from its original position and establishes the future anterior end of the larva. A number of mesodermal cells (anterior mesoderm) delaminate from the anterior endodermal-ectodermal boundary (Figure 2I and 3C). At early larva stage (40hpf), the former blastopore is located anterior-ventrally, where it eventually forms the future mouth of the larva. The archenteron then narrows and becomes a posteriorly blind tube. The ectoderm grows massively and forms a pre-oral lobe, which protrudes anteriorly and ventrally of the mouth. Some mesodermal cells spread into the pre-oral lobe and others migrate posteriorly to form two lateral tiers along both sides of the archenteron. The posterior-ventral region of ectoderm thickens and leads to the formation of the tentacular ridge; which will later give rise to the first pair of tentacles (Figure 2J and 3D). At 50-60hpf, the pre-tentacle actinotrocha larva is almost formed. The pre-oral lobe becomes more prominent. The archenteron differentiates into esophagus (foregut), stomach (midgut) and intestine (hindgut) and the anus opens after the junction of intestinal and ectodermal cells. The tentacle bulbs are evident and the protonephridial primordia are established (Figure 2K and 3E). At 100hpf (5days), the larva has already 3 pairs of tentacles and a well-defined telotroch around the anus (Figure 3F). The posterior mesoderm forms at the junction of the stomach and the intestine, the protonephridia are evident and the mid part of the stomach protrudes to develop a stomach diverticulum (Figure 3F).

**Figure 3.**
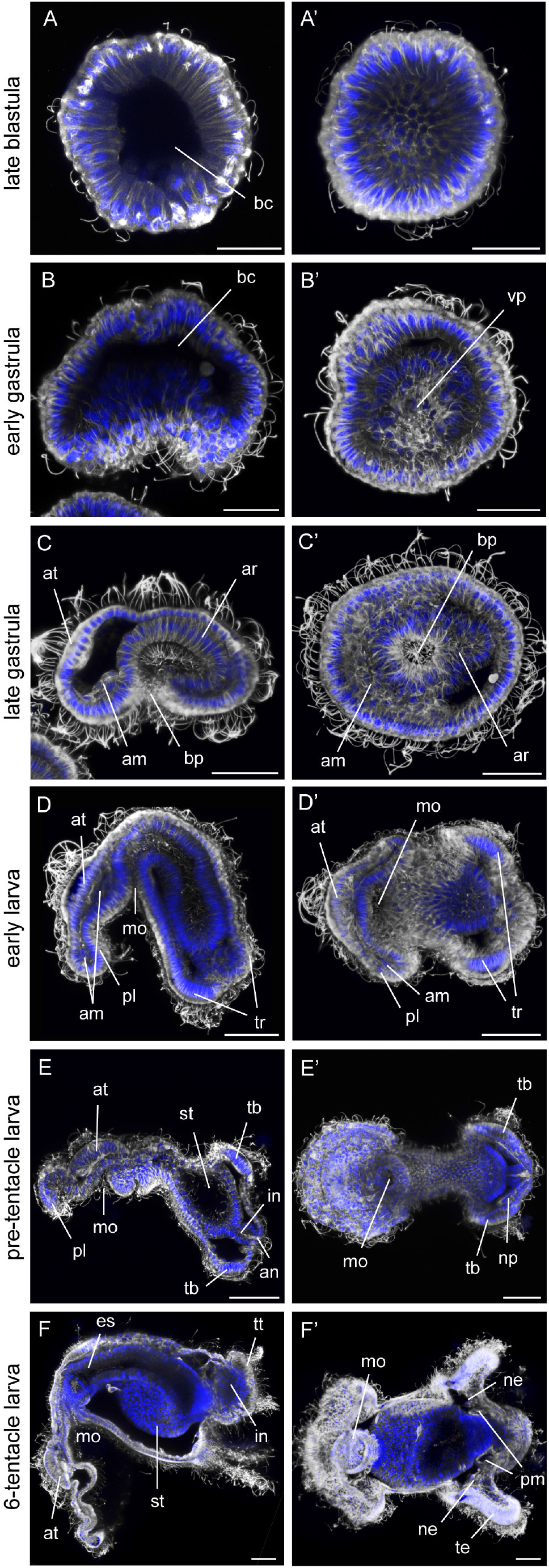
Immunohistochemistry on blastula, gastrula and larva stages of *P. harmeri*. Immunohistochemistry on blastula, gastrula and larva stages labeled against acetylated tubulin (grey) and DAPI (blue). Panels B-F depict embryos in lateral view (Iv) and panels B’-F’ show embryos in vegetal view (vv). In panels depicting gastrulae and larvae stages, anterior is to the left, am, anterior mesoderm; an, anus; ar, archenteron; at, apical organ; be, blastocoel; bp, blastopore; es, esophagus; in, intestine; mo, mouth; ne, nephridium; np, nephridial primordium; pl, pre-oral lobe; pm, posterior mesoderm; st, stomach; tb, tentacle bulb; te, tentacle; tr, tentacular ridge; tt, telotroch; vp, vegetal plate. Scale bar: 25 μm.

### Molecular patterning of the endomesoderm of *P. harmeri*

To reveal the spatial and temporal appearance of endodermal and mesodermal fates, we analyzed the expression of evolutionarily conserved molecular markers associated with the development of endomesodermal tissues, *foxA*, *gata4/5/6* and *twist*, in blastula, gastrula and larva stages of *P. harmeri*.

*FoxA* is already expressed at the blastula stage, in few cells of the vegetal pole (Figure 4A and Figure 5A). At the early gastrula stage, the gene is expressed asymmetrically in the anterior ventrolateral ectoderm and in the whole vegetal plate (Figure 4B and Figure 5D). Later, at the late gastrula stage, the expression of *foxA* is retained mostly around the blastopore and faintly in the invaginating archenteron (Figure 4C). In the early larva, *foxA* expression is seen around the mouth, and the ventral ectoderm (Figure 4D), overlapping with *bra*, where it remains at the pre-tentacle and 6-tentacle larva stages (Figure 4E-F and Figure 5J).

**Figure 4.**
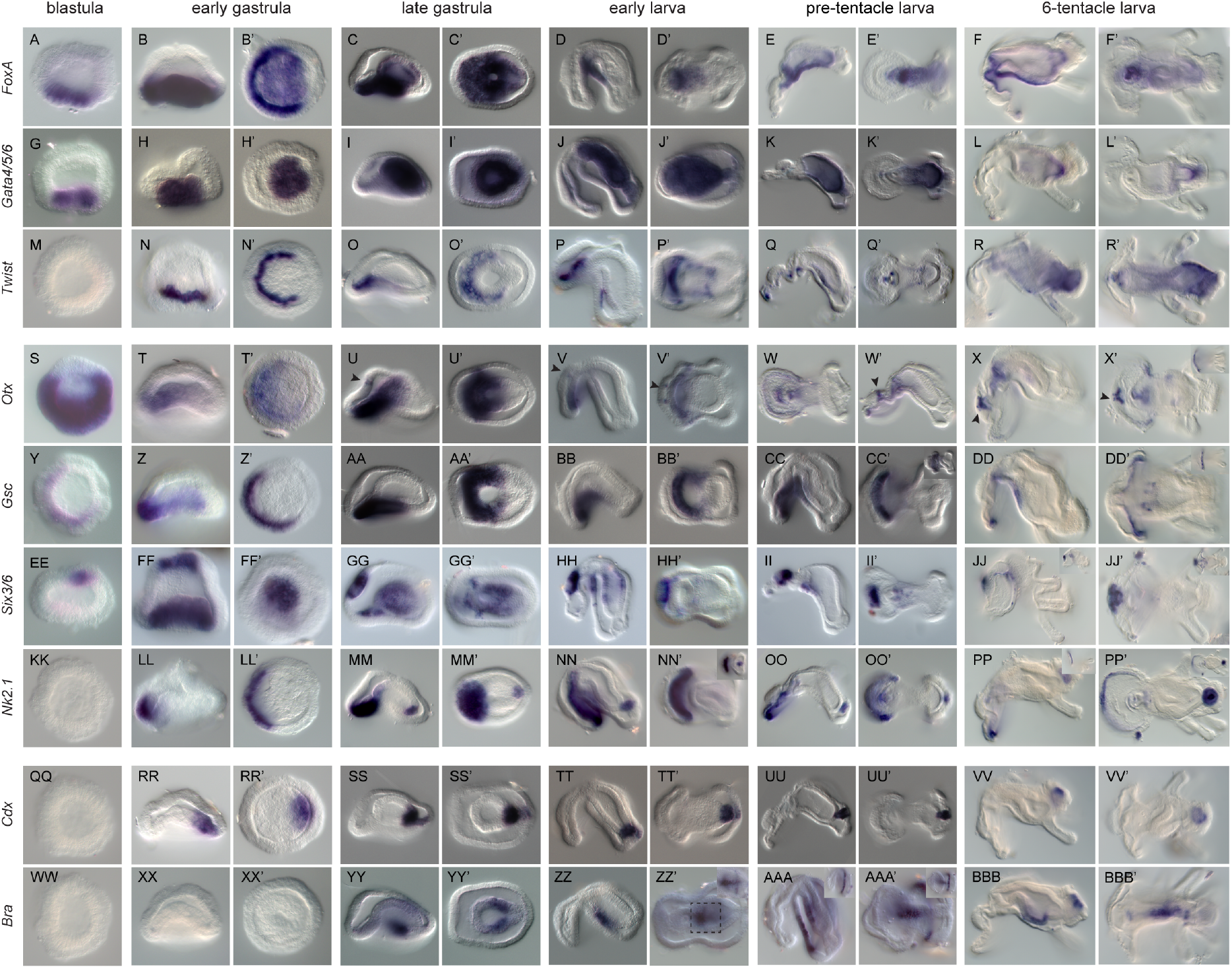
Expression of endomesodermal, anterior and posterior markers during the embryonic development of *P. harmeri*. WMISH of *otx, gsc, six3/6, nk2.1, cdx, bra, foxA, gata4/5/6* and *twist* in blastula, early gastrulae, late gastrulae, early larvae, pre-tentacle larvae and 6-tentacle larvae of *P. harmeri*. The panels of the first columns **(A-BBB)** depict embryos in lateral view and the panels of the second columns show embryos in vegetal view **(A’-BBB’).** Insets in panels X’, CC’, DD’, JJ-JJ’, NN’, PP-PP’ and AAA-AAA’ show different focal planes of the embryos. Black arrow indicates the ectodermal expression of *otx* at the domain that gives rise to the apical organ. The inset in panel ZZ’ shows a higher magnification of the indicated domain. In panels depicting gastrulae and larvae stages, anterior is to the left.

*Gata4/5/6* is expressed at the blastula stage, in few cells of the vegetal pole, overlapping with *foxA* (Figure 4G and Figure 5C). At the early gastrula stage, *gata4/5/6* is expressed in the vegetal plate that will later ingress to form the archenteron (Figure 4H). At the late gastrula stage, transcripts of the gene are only detected in the invaginating archenteron (Figure 4I), where they remain at the early, pre-tentacle and 6-tentacle larva stages (Figure 4J-K). At the 6-tentacle larva stage the expression of *gata4/5/6* is restricted to the pyloric sphincter (Figure 4L).

The mesodermal marker *twist* starts to be expressed at the early gastrula stage, in an anterior ventrolateral cell population of the vegetal plate, located adjacently to the expression of *foxA* (Figure 4N and Figure 5D). At the late gastrula stage, *twist* expression is detected at the anterior mesoderm, where it overlaps with *six3/6* expression (Figure 4O and Figure 5H). As some of these anterior mesodermal cells migrate posteriorly, forming two lateral tiers along both sides of the archenteron, at the early larva stage *twist* is expressed in both the pre-oral mesoderm and these two ventro-lateral tiers (Figure 4P). In the pre-tentacle larva, the expression of *twist* remains in clusters of cells of the pre-oral and post-oral mesoderm, and the two ventro-lateral tiers of mesoderm (Figure 4Q). At the 6-tentacle larva stage, *twist* expression is additionally seen at the posterior and tentacular mesoderm (Figure 4R).

**Figure 5:**
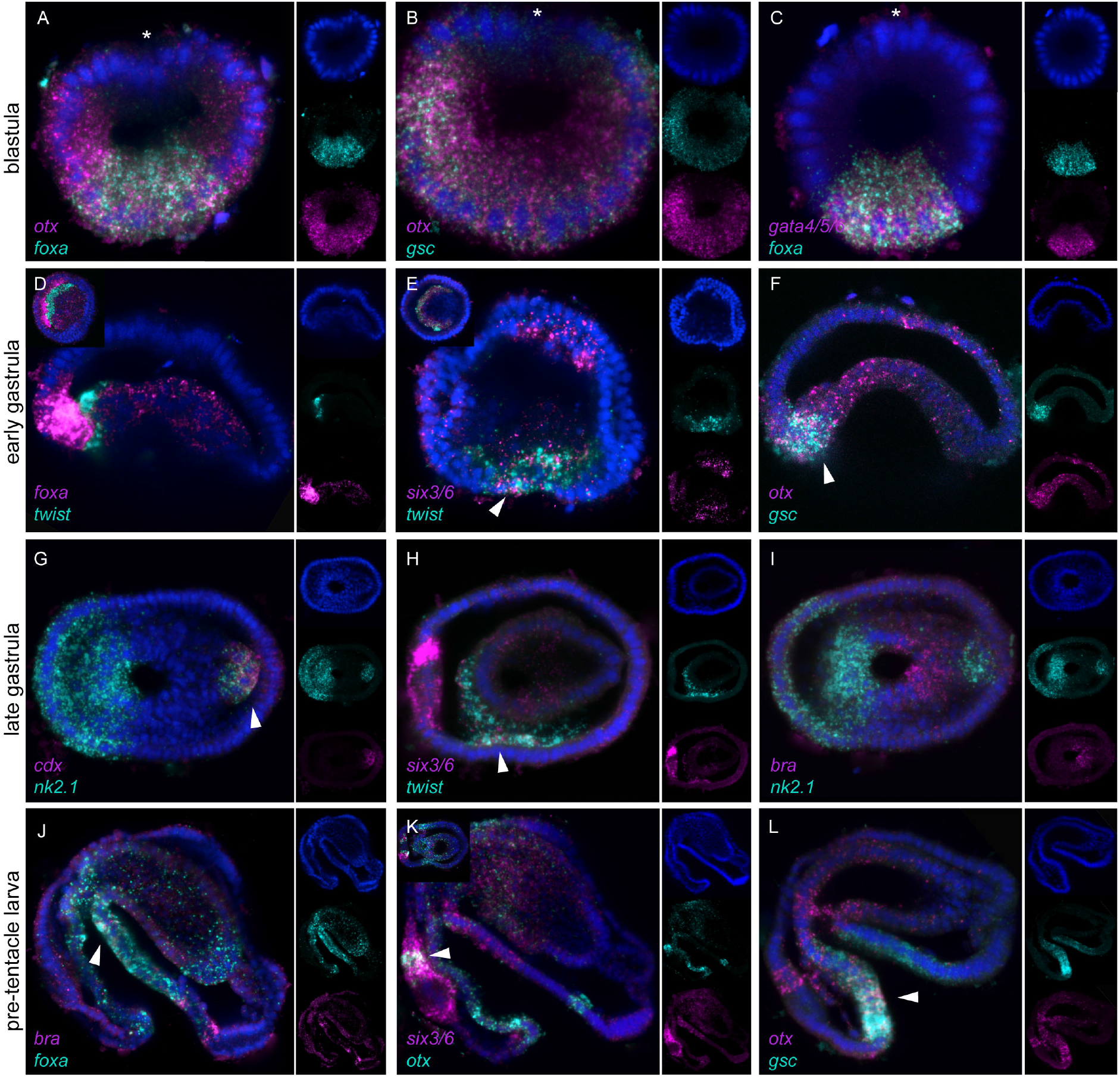
Co-expression analysis of marker genes by double fluorescent WMISH. Relative spatial expression of *otx* and *foxA* **(A),** *otx* and *gsc* **(B, F, L)**, *gata4/5/6* and *foxA* **(C)**, *foxA* and *twist* **(D)**, *six3/6* and *twist cdx* and *nk2.1* **(G),** *bra* and *nk2.1* (I), *bra* and *foxA* (J) and *six3/6* and *otx* **(K)**. Right insets in panels D, E and K show embryos in vegetal view. Every picture is a full projection ofmerged confocal stacks. Nuclei are stained blue with DAPI. Anterior is to the left.

### Anterior-posterior molecular patterning of *P. harmeri*

To identify the segregation of the embryonic fates along the anterior-posterior axis, we analyzed the expression of genes with a conserved anterior expression, such as *orthodenticle (otx), goosecoid (gsc), six3/6* and *nk2.1*, and genes commonly involved in the specification of posterior tissues, such as *caudal (cdx)* and *brachyury (bra)*, in blastula, gastrula and larva stages of *P. harmeri*.

*Otx* is expressed broadly at the blastula stage, throughout the vegetal hemisphere in to the animal hemisphere, but excluding the animal pole (Figure 4S and Figure 5A-B), but at the early gastrula stage, the gene is restricted in the anterior lip of the blastopore and the anterior part of the invaginating archenteron (Figure 4T and Figure 5F). In the late gastrula, the expression of the gene remains in the anterior blastoporal lip and the anterior domain of the invaginating archenteron, and also initiates in few cells of the anterior ectoderm, a region that will form the future apical organ (Figure 4U). At the early larva stage, the gene is expressed in the most anterior region of the ventral ectoderm of the pre-oral lobe, that will later form the neuronal-rich edge of the preoral hood, the mouth, and two cell clusters of the apical organ (Figure 4V), where it remains at the pre-tentacle and 6-tentacle larva stages (Figure 4W-X and Figure 5L). Additionally, *otx* expression is detected in a small cell cluster of the most posterior ventral ectoderm (Figure 5K).

*Gsc* expression initiates on one side of the blastula, within the *otx*-positive domain (Figure 4Y and Figure 5B). Later, at the early gastrula stage, *gsc* is expressed in an ectodermal domain of the vegetal plate that corresponds to the anterior blastoporal lip (Figure 4Z), overlapping with *otx* at the most anterior part (Figure 5F). At the late gastrula stage, the gene remains active around the blastopore (Figure 4AA). In the early larva, the expression of *gsc* is restricted in the ventral ectoderm of the pre-oral lobe and the mouth, overlapping with *otx* (Figure 4BB), where it remains at the pre-tentacle and 6-tentacle larva stages (Figure 4CC-DD and Figure 5L). *Six3/6* is expressed in approximate 4-5 cells of the animal pole already at the blastula stage (Figure 4EE). At the early gastrula stage, the expression of the gene remains in the animal pole, in the region that will give rise to the future apical organ, and an anterior ventrolateral cell population of the vegetal plate, which corresponds to the anterior mesoderm (Figure 4FF and Figure 5E). In the late gastrula, transcripts of the gene are found at the apical organ, the anterior mesoderm and a few scattered cells of the archenteron (Figure 4GG and Figure 5H). At the early larva stage, *six3/6* expression is seen in the apical organ, overlapping partially with *otx*, clusters of cells of the ventral ectoderm and the developing midgut (Figure 4HH), and some pre-oral mesodermal cells, where it remains at the pre-tentacle and 6-tentacle larva stages (Figure 4II-JJ and Figure 5K). At the 6-tentacle larva stage, transcripts of *six3/6* are also detected in individual cells of the edge of the preoral hood, possibly muscles (Figure 4JJ). *Nk2.1* is first expressed in an ectodermal domain of the vegetal plate, where the anterior blastoporal lip will form, at the early gastrula stage (Figure 4LL). At the late gastrula stage, the expression of the gene remains in the anterior blastoporal lip and also initiates in the most posterior region of the developing archenteron (Figure 4MM and Figure 5G, I). At the early larva stage, *nk2.1* is expressed at the ventral ectoderm of the pre-oral lobe and the intestine (Figure 4NN), where it remains at the pre-tentacle and 6-tentacle larva stages (Figure 4OO-PP).

*Cdx* starts to be expressed at the early gastrula stage, in one group of cells of the vegetal plate that correspond to the posterior blastoporal lip (Figure 4RR). In the late gastrula, the expression of the gene in detected in the posterior region of the developing archenteron, where it overlaps with *nk2.1*, and in the posterior ectoderm that will later form the anus (Figure 4SS and Figure 5G). At the early, pre-tentacle and 6-tentacle larva stages, *cdx* expression is restricted to the intestine (Figure 4TT-VV).

The expression of *bra* initiates only at the late gastrula stage, in the posterior blastoporal lip that will give rise to the developing midgut (Figure 4YY and Figure 5I). At the early larva stage, the expression of the gene shifts to the ventral domain of the midgut and few cells of the posterior ventral ectoderm (Figure 4ZZ). At the pre-tentacle larva stage transcripts of *bra* are retained in the ventral midgut and expand in more cells of the ventral ectoderm and the posterior ciliary band (Figure 4AAA and Figure 5J). In the 6-tentacle larva, transcripts of *bra* are detected also in the most posterior domain of the intestine, similarly to *cdx*, as well as the ventral ectoderm and the stomach diverticulum (Figure 4BBB).

A summary of all gene expression patterns described in this study is provided in Figure 6.

**Figure 6.**
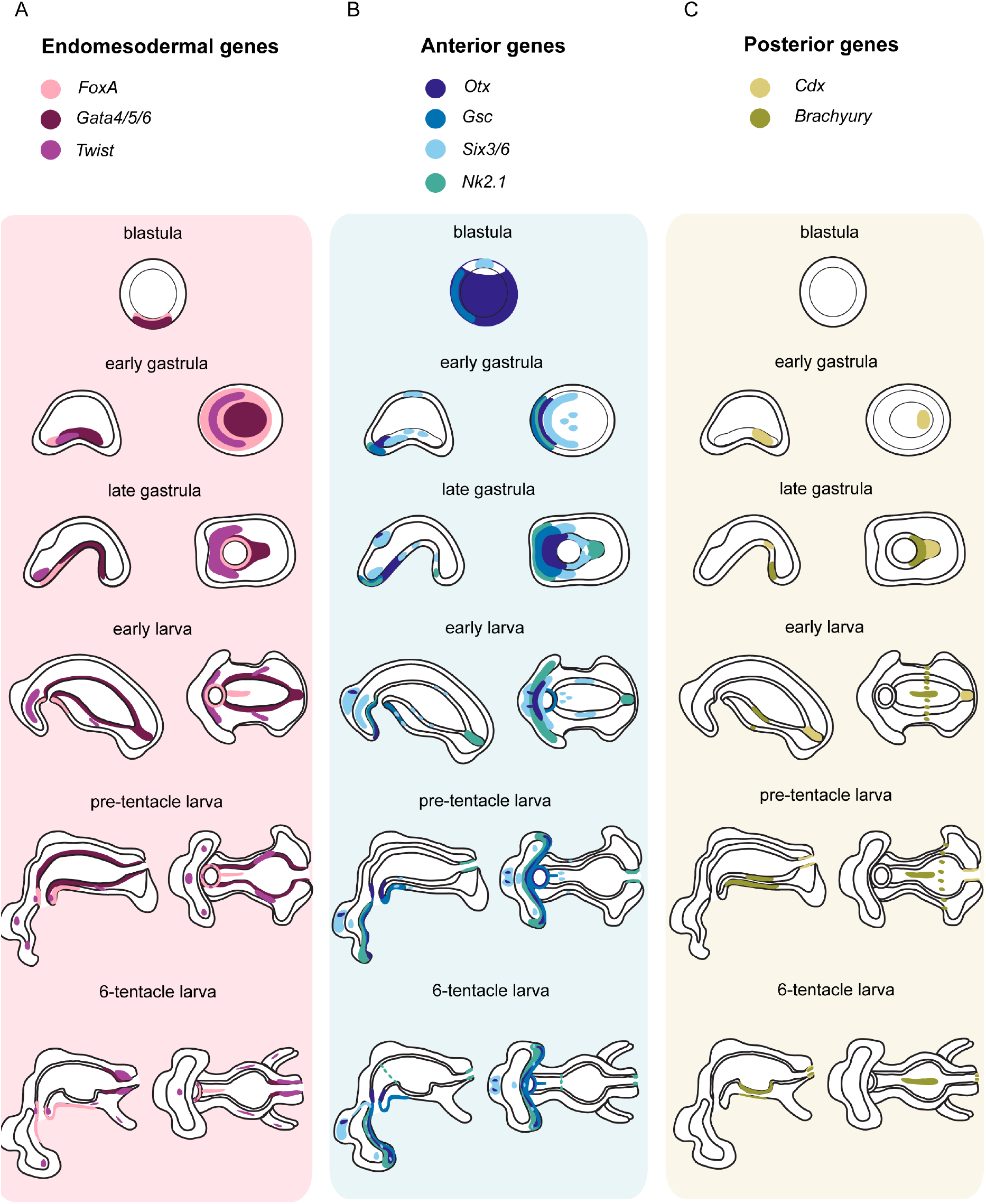
Summary of gene expression during *P. harmeri* embryonic development. Schematic representation of the expression patterns of endomesodermal, anterior and posterior markers during embryonic development of *P. harmeri*. (**A**) The endodermal genes *foxA* and *gata4/5/6* are expressed in the vegetal plate in blastula and later on are patterning the formation of the archenteron. *FoxA* eventually confines in the foregut, whilst *gata4/5/6* is expressed in the midgut. The mesodermal marker *twist* is labeling the anterior and posterior mesoderm and its derivatives. (**B**) The anterior gene *six3/6* is expressed in the animal pole in blastula and at the gastrula stage is also activated in the anterior mesoderm and clusters of cells of the future midgut. At the early, pre-tentacle and 6-tentacle larva stages *six3/6* is restricted in the apical organ, anterior mesoderm and the oral ectoderm. *Otx* is expressed broadly at the blastula stage, and in gastrula it labels the anterior lip of the blastopore, adjacent to the expression of *nk2.1* and *gsc*. At the gastrula stage, *otx, nk2.1* and *gsc* are labeling the anterior-ventral ectoderm. *Otx* is also expressed in the future apical organ and the future midgut, and *nk2*.1 is additionally labeling the future hindgut. Later on, *otx* and *nk2.1* are marking the ventral ectoderm of the pre-oral lobe. *Otx* together with *gsc* are demarcating the mouth. Additionally, *otx* labels the apical organ and *nk2.1* is expressed strongly in the intestine and in the cardiac sphincter. (**C**) The posterior markers *bra* and *cdx* are expressed in the posterior lip of the blastopore at the gastrula stage. *Bra* is expressed in the ventral midgut, the ventral ectoderm and the posterior ciliary band, whilst *cdx* confines in the intestine. At the 6-tentacle larva stage *bra* is activated in the intestine, the stomach diverticulum and the ventral ectoderm.

## Discussion

### Novelties and evolutionary dynamics in marker gene expression

Comparisons of embryos from different evolutionary lineages have shown that the molecular interplay of axial and cellular specification is characterized by remarkable conservation of expression patterns for many genes, but also by important lineage specific novelties [46–50], To better understand the ancestral molecular underpinnings of cellular identities, more molecular data are needed from understudied embryos such as the phoronids. Here, we analyzed the expression patterns of the evolutionarily conserved anterior (*otx, gsc, six3/6, nk2.1*), posterior (*cdx, bra*) and endomesodermal (*foxA, gata4/5/6, twist*) markers in the phoronid *P. harmeri*. Our study shows an overall conservation in embryonic molecular patterning, but also highlights a number of unexpected expression profiles. For example, *foxA*, *gata4/5/6* and *cdx* show conserved expression in patterning the development and regionalization of the phoronid embryonic gut, with *foxA* expressed in the presumptive foregut, *gata4/5/6* demarcating the midgut and *cdx* confined to the hindgut, similar to what is reported in ecdysozoans [41, 51, 52], echinoderms [42, 53, 54], spiralians [55–57], and vertebrates [58–60], Another gene that labels the hindgut in phoronids is *nk2.1*, similarly to what is reported in some annelids [40], acoels [61], hemichordates [62], and cephalochordates [63], Surprisingly, *brachyury*, an evolutionary conserved transcription factor often associated with gastrulation movements and patterning of the mouth and hindgut [45, 52, 64–66], seems to be unrelated with gastrulation and mouth patterning in phoronids. *Bra* starts to be expressed only after gastrulation initiates, and exhibits a dynamic expression pattern, labeling not only the hindgut, but also the ventral region of the midgut, the ventral ectoderm, and the posterior ciliary band. Expression of *bra* in the ventral ectoderm has also been reported in acoels [61], and expression in the developing ciliated band (velar rudiment) has been reported in the mollusc *Crepidula* [67], However, it is not clear whether this ‘module’ of *bra* expression is conserved within these taxa. Interestingly, *six3/6*, a well-conserved anterior marker [34, 36, 37, 61, 68–70], also shows remarkably dynamic expression in phoronids, demarcating not only the apical organ and the oral ectoderm, but also clusters of cells of the developing midgut and the anterior mesoderm. Similar endomesodermal domains of expression have been reported in other taxa too; for instance *six3/6* is expressed in the endomesoderm of brachiopods and bryozoans [69, 71], hemichordates [34], the mesenchyme cells of echinoderms [72] and the endoderm of cnidarians [68], Finally, the expression of the evolutionary conserved anterior/CNS marker *nk2.1* [34, 36, 40, 67, 73] displays a notable difference in the phoronid compared to other animals examined, as this gene is not expressed in the future anterior end of the larva, where the apical organ will form, but rather is localized to the edge of the preoral hood, likely in developing neurons. Overall, our observations allowed us to pinpoint similarities in the embryonic molecular patterning of phoronids in relation to other animals, but also identify interesting differences.

### Comparative molecular embryology between Phoronida and Brachiopoda

Previous embryonic comparisons based on fate maps between a number of brachiopod species (*T. transversa, Hemithiris* sp., *Terebratulina* sp. and *N. anomala*) and phoronids (*Phoronis vancouverensis*), suggested differences in the timing of axis and regional specification [12, 74–77], For instance, in *P. vancouverensis, T. transversa, Hemithiris* sp. and *Terebratulina* sp., axis formation is related to the movement of cells along the dorsal side of the future anterior–posterior axis of the larva during late gastrulation, whilst in *N. anomala* the larval anterior–posterior axis corresponds to the animal-vegetal axis of the egg and that axis is set up already before the blastula stage [12, 74–77], Recent molecular data from *T. transversa* and *N. anomala* development also support the notion that an anterior-posterior molecular re-patterning of the blastopore occurs at the gastrula stage in *T. transversa*, which takes place before axial elongation, unlike in *N.anomala*, where such a symmetry-breaking event is absent [71].

Our molecular comparison of phoronid and brachiopod development confirmed some of the conclusions from the aforementioned studies and revealed some conservation of molecular patterning and gene topology between *P. harmeri*, *T. transversa* and *N. anomala*, but also showed differences related to their different developmental modes (Figure 7). In particular, the endomesodermal fates of both brachiopod species and *P. harmeri* are similar in that the vegetal pole of the blastula is the site where endoderm and mesoderm originate, as revealed by the expression of *foxA, twist* and *gata4/5/6* endomesodermal markers ([43, 71], this study). However, a notable difference was observed in the patterning of phoronid mesoderm. In *P. harmeri*, the molecular specification of anterior mesoderm occurs at the early gastrula stage, as revealed from the expression of the mesodermal marker *twist;* thus the timing of mesoderm specification is more similar to *N. anomala* than to *T. transversa*, where mesoderm is already specified at the blastula stage [43, 71], In addition to *twist*, mesoderm development in phoronids is also patterned by *six3/6*, a conserved anterior marker, in contrast to brachiopods, where *six3/6* is solely expressed in the anterior ectoderm and endoderm [36, 71]. However, no expression of *six3/6* is observed in the posterior mesoderm of phoronids, at the 6-tentacle larva stage. This difference in mesodermal patterning might be related to different embryological sources of the anterior and posterior mesoderm [13, 20], More molecular studies on mesoderm development are needed in phoronids, to clarify whether the formation of posterior mesoderm utilizes different molecular mechanisms from anterior mesoderm.

**Figure 7.**
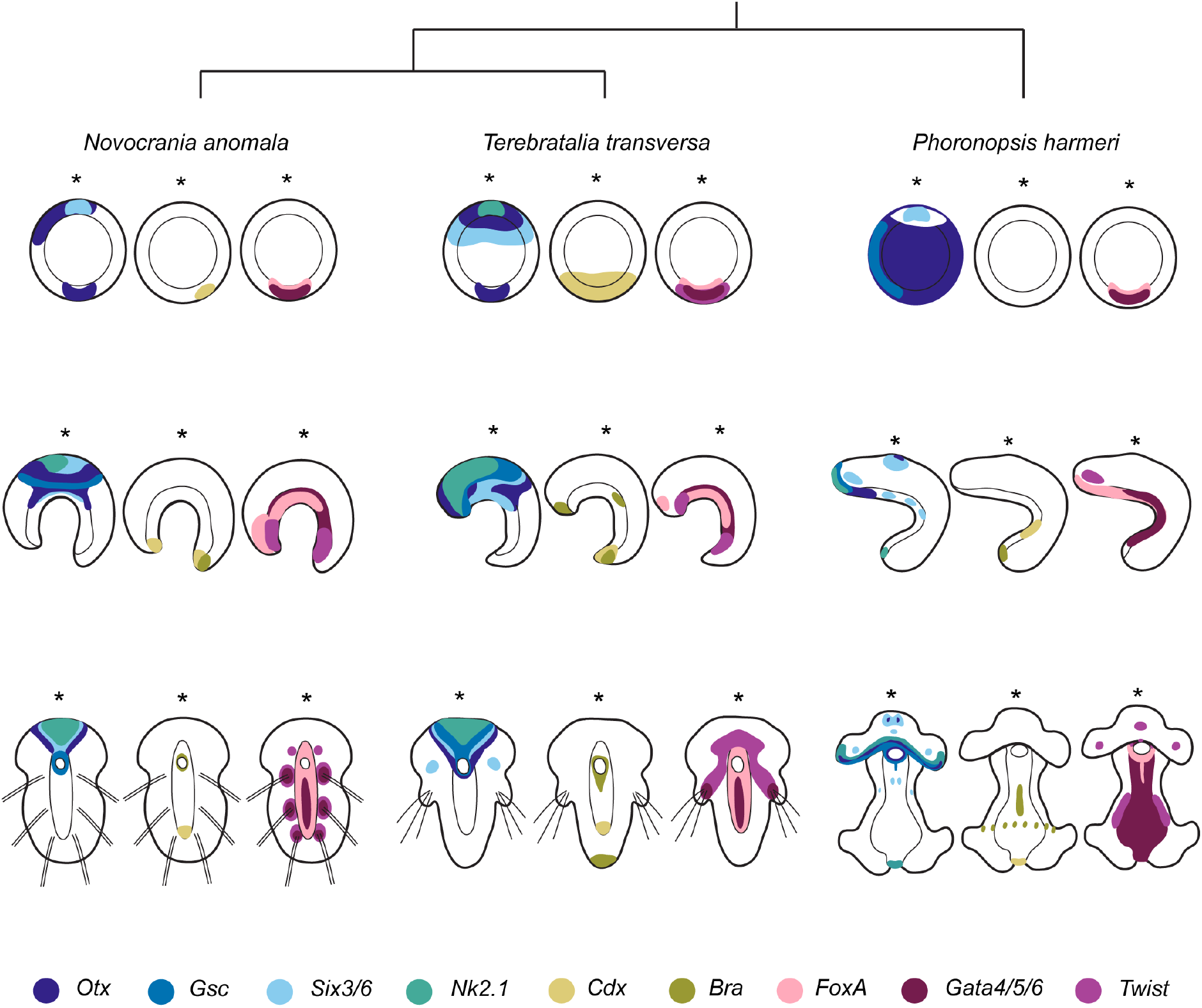
Comparison of embryonic gene expression patterns in representative developmental stages of *Novocrania anomala, Terebratalia transversa* and *Phoronopsis harmeri*. Schematic representation of the expression patterns of endomesodermal, anterior and posterior markers in blastula, mid gastrula and larva stages of two members of Brachiopoda (*N. anomala* and *T. transversa*) and one member of Phoronida (*P. harmeri*). The asterisk is indicating the anterior domain.

Other observed differences relate to the specification of posterior tissues. In *P. harmeri* posterior fates seem to be not yet established at the blastula stage, as indicated by the lack of expression of the posterior marker *cdx*, in contrast to brachiopods, where *cdx* is localized at the vegetal pole of the blastula and already demarcates the future posterior territory of the embryo [71].

In *P. harmeri, cdx* starts to be expressed at early gastrula stage only in the posterior blastoporal lip, similarly to *T.transversa* but not to *N.anomala*, where the gene remains activated around the blastopore till early larva stage [71], The restriction of *cdx* in the posterior blastoporal lip in *P. harmeri* is related to the different gastrulation mode and blastoporal fate observed between species (*P. harmeri* and *T. transversa* exhibit protostomy, while *N. anomala* is deuterostomic). Nevertheless, in all three species the expression of *cdx* will eventually be restricted to the posterior region of the larval gut (that corresponds to the intestine in *P. harmeri*) (this study, [71]). One more gene associated with posterior identity and mouth formation in brachiopods [71], *bra*, exhibits a unique pattern of expression in *P. harmeri*, since transcripts of the gene are found in the ventral ectoderm and the ventral region of the developing midgut that will form the stomach diverticulum, a distinct structure of the midgut rich in secretory cells with enormous endoplasmic reticulum [78]. Other interesting properties of *bra* expression pattern are the late activation of the gene in the most posterior part of the intestine and the absence of expression in the mouth. The expression of *bra* in the intestine of the phoronid 6-tentacle larva stage might be related to the fact that during metamorphosis the larval intestine is kept and transforms into the intestine of the juvenile [15, 78].

Another intriguing difference is the potential role of *nk2.1* in patterning posterior tissues in *P. harmeri*, whilst in brachiopods the orthologous gene is only involved in the specification of the anterior structures [36, 71]. The role of *nk2.1* in posterior patterning and hindgut formation in phoronids, but not in brachiopods, might be attributed to the fact that *P. harmeri* possess a planktotrophic larva with a tripartite, functional gut, whilst *T. transversa* and *N. anomala* form lecithotrophic larva with a blind gut, lacking a hindgut. Unfortunately, expression data are not available for the planktotrophic larva of linguliform brachiopods, which would elucidate whether *nk2.1* has a conserved role in patterning the hindgut of both phoronids and brachiopods, or this expression has been co-opted in phoronids.

Differences are also seen regarding the relative position of the future anterior structures and oral ectoderm patterning during gastrulation between phoronids and brachiopods. In all three species the future anterior end of the larva seems to be specified by blastula stage, as seen from the expression of the anterior marker, *six3/6* ([36, 71], this study). However, neither *nk2.1* nor *gsc* demarcate the future anterior end of *P. harmeri* larva, as described in brachiopods ([36, 71], this study). The absence of expression of these anterior markers in the future anterior end of *P. harmeri* likely reflects the uncoupling of the animal-vegetal and anterior-posterior axes observed during phoronid development [12, 14, 16, 19, 20, 22, 24, 28]. *Nk2.1* and *gsc* are exclusively expressed in the anterior lip of the blastopore in *P. harmeri* during gastrulation, similarly to what is reported in *T. transversa*, but different from *N. anomala*, where the expression of these genes is seen mainly in the anterior region of the embryo that remains separated from the blastopore throughout development (this study, [71]). The expression of anterior markers in the anterior blastoporal lip in *P. harmeri* suggests a contribution of this region to the oral ectoderm (and mouth) formation. This is similar to what is reported in the protostomic *T. transversa* [71], and therefore reflects a common molecular patterning during gastrulation due to the protostomic mode of development in both organisms.

In general, the developmental patterning and the cellular specification of the rhynchonelliform brachiopod *T. transversa* and *P. harmeri* appear to be more similar than the craniiform brachiopod *N. anomala*, with the exception of the timing of mesoderm specification and mesodermal patterning. Similar conclusions emerged from the previous comparative embryonic fate map studies conducted between brachiopods and phoronids [12, 74–77], According to these studies, phoronids, rhynchonelliform and linguliform brachiopods share identical fate maps, suggesting that the last common ancestor of brachiopods and phoronids likely shared an early molecular embryonic patterning similar to the extant rhynchonelliform brachiopods and phoronids.

## Conclusions

In this work, we provide a molecular characterization of the embryogenesis of the phoronid *P. harmeri*, with detailed gene expression profiling of marker genes related to cell and axis specification during animal development. We show that the future endodermal and anterior territories appear to be specified by the blastula stage, in contrast to posterior fates that are established later in development. Comparing the embryonic patterning of *P. harmeri* with available data of brachiopods, the proposed sister group to Phoronida, we observe more similarities with rhynchonelliform than with craniiform brachiopods, which is likely related to their different gastrulation modes. Taken together, our findings illustrate how variations in the early molecular pattering can result in differences in axis and cellular specification, and present a basis for future studies on the evolution of the development of Lophophorata.

## Methods

### Animal systems

Adult specimens of *Phoronopsis harmeri* Pixell, 1912 were collected at the sand flat of Gaffney point, close to the main channel, at lowtide, in Bodega bay, California, USA(38° 18’ 51.9012” N 123° 3’ 12.3012” W), in April. Eggs were obtained from female animals by puncturing the posterior body wall. Fertilization occurred instantly, the moment of the sacrifice. The embryos were kept in clean seawater at 9°C and were fed with concentrated Rhodomonas algae from the pre-tentacle larva stage onwards.

### Gene cloning and orthology assignment

Putative orthologous sequences of genes of interest were identified by tBLASTx search against the transcriptome of *Phoronopsis harmeri*. Gene orthology of genes of interest identified by tBLASTx was tested by reciprocal BLAST against NCBI Genbank and followed by phylogenetic analyses. Amino acid alignments were made with MUSCLE. RAxML (version 8.2.9) was used to conduct a maximum likelihood phylogenetic analysis. Fragments of the genes of interest were amplified from cDNA of *P. harmeri* by PCR using gene specific primers. PCR products were purified and cloned into a pGEM-T Easy vector (Promega, USA) according to the manufacturer’s instruction and the identity of inserts confirmed by sequencing.

### Data Availability

All newly determined sequences have been deposited in GenBank under accession numbers MN431422 - MN431430. Primer sequences are available on request.

### Whole Mount *In Situ* Hybridization

Embryos were manually collected, fixed and processed for *in situ* hybridization as described in [61]. Labeled antisense RNA probes were transcribed from linearized DNA using digoxigenin-ll-UTP (Roche, USA) according to the manufacturer’s instructions.

### Whole Mount Immunohistochemistry

Animals were collected manually, fixed in 4% paraformaldehyde in SW for 60 minutes, washed 3 times in PBT and incubated in 4% sheep serum in PBT for 30 min. The animals were then incubated with commercially available primary antibodies (anti-acetylated and anti-tyrosinated tubulin mouse monoclonal antibody, dilution 1:250 (Sigma-Aldrich, USA) overnight at 4°C, washed 10 times in PBT, and followed by incubation in 4% sheep serum in PBT for 30 min. Specimens were then incubated with a secondary antibody overnight at 4°C followed by 5 washes in PTW. Nuclei were stained with DAPI.

### Documentation

Colorimetric WMISH specimens were imaged with a Zeiss AxioCam HRc mounted on a Zeiss Axioscope A1 equipped with Nomarski optics and processed through Photoshop CS6 (Adobe). Fluorescent-labeled specimens were analyzed with a SP5 confocal laser microscope (Leica, Germany) and processed by the ImageJ software version 2.0.0-rc-42/1.50d (Wayne Rasband, NIH). Figure plates were arranged with Illustrator CS6 (Adobe).

## Acknowledgements

We thank Andrea Orus Alcalde and Ludwik Gasiorowski from Hejnol’s group for contributing with animal collection and spawning. Kevin Uhlinger for help with rearing larvae and Karl Menard from Bodega Bay Marine Lab for help with collecting adults. We thank Chema Martin-Duran for reading the manuscript.

## Authors’ contributions

CA, CL, MQM and AH designed the study. CL made and provided the transcriptome. YP and CA performed the collections and CA conducted the experiments. CA, CL, MQM and AH analyzed the data. CA and AH wrote the manuscript. All authors read and approved the final manuscript.

---

**Supplementary Figure 1: Orthology analysis.** Putative orthologous sequences of genes of interest were identified by tBLASTx search against the transcriptome of *P. harmeri*. Bayesian phylogenetic analysis is supporting orthology. Names of genes or proteins, if available, follow the name of organism(s). *P. harmeri* sequences are highlighted in red. Ph, *Phoronopsis harmeri;* Na, *Novocrania anomala;* Tt, *Terebratalia transversa;* Mm, *Membranipora membranacea;* Hs, *Homo sapiens;* Xl, *Xenopus laevis;* Xt, *Xenopus tropicalis;* Dr, *Danio rerio;* Mm, *mus musculus;* Gg, *Gallu gallus;* Sk, *Saccoglossus kowalevskii;* Pf, *Ptychodera flava;* Sp, *Strongylocentrotus purpuratus;* Pl, *Paracentrotus lividus;* Lv, *Lytechinus variegatus;* Am, *Asterina miniata;* At, *Archaster typicus;* Ci, *Ciona intestinalis;* HI, *Halocynthia roretzi;* Od, *Oikopleura dioica;* Bf, *Branchiostoma floridae;* Sm, *Strigamia maritima;* Dm, *Drosophila melanogaster;* Tc, *Tribolium castaneum;* Lg, *Lottia gigantea; Euprymna;* Cf, *Crepidula fornicata;* Ml, *Macrostomum lignano;* Sm, *Schmidtea mediterranean* Spoly, *Schmidtea polychroa;* Pv, *Prostheceraeus vittatus;* Gt, *Girardia tigrina;* Ct, *Capitella teleta;* Of, *Owenia fusiformis;* Pd, *Platynereis dumerilii;* He, *Hydroides elegans;* Tt, *Tubifex tubifex; Chaetopterus; Phascolion;* Ap, *Apis mellifera;* Pc, *Priapulus caudatus; Achaearanea;* Bm, *Bombyx mori;* Ha, *Helicoverpa armigera;* Nv, *Nematostella vectensis; Hydractinia; Hydra;* Cr, *Cladonema radiatum;* Aa, *Aurelia aurita;* Pc, *Podocoryna carnea;* Ms, *Meara stichopi;* Cm, *Convolutriloba macropyga;* Ta, *Trichoplax adhaerens;* Sc, *Sycon ciliatum;* Pb, *Pleurobrachia bachei; Diplosoma*.

