## Supplementary figures and images for "Molecular patterning during the development of *Phoronopsis harmeri* reveals similarities to rhynchonelliform brachiopods"

### Supplementary Information

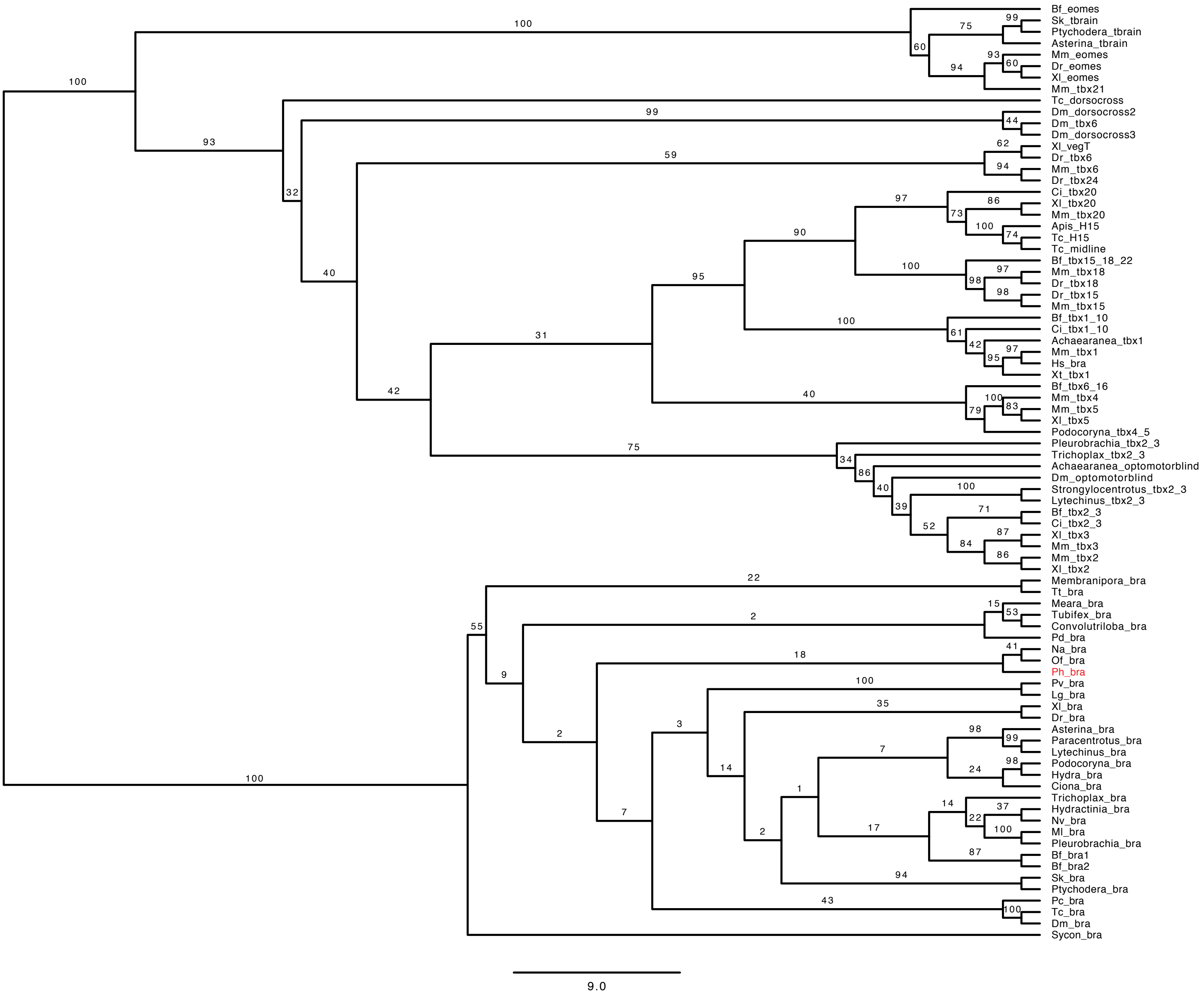

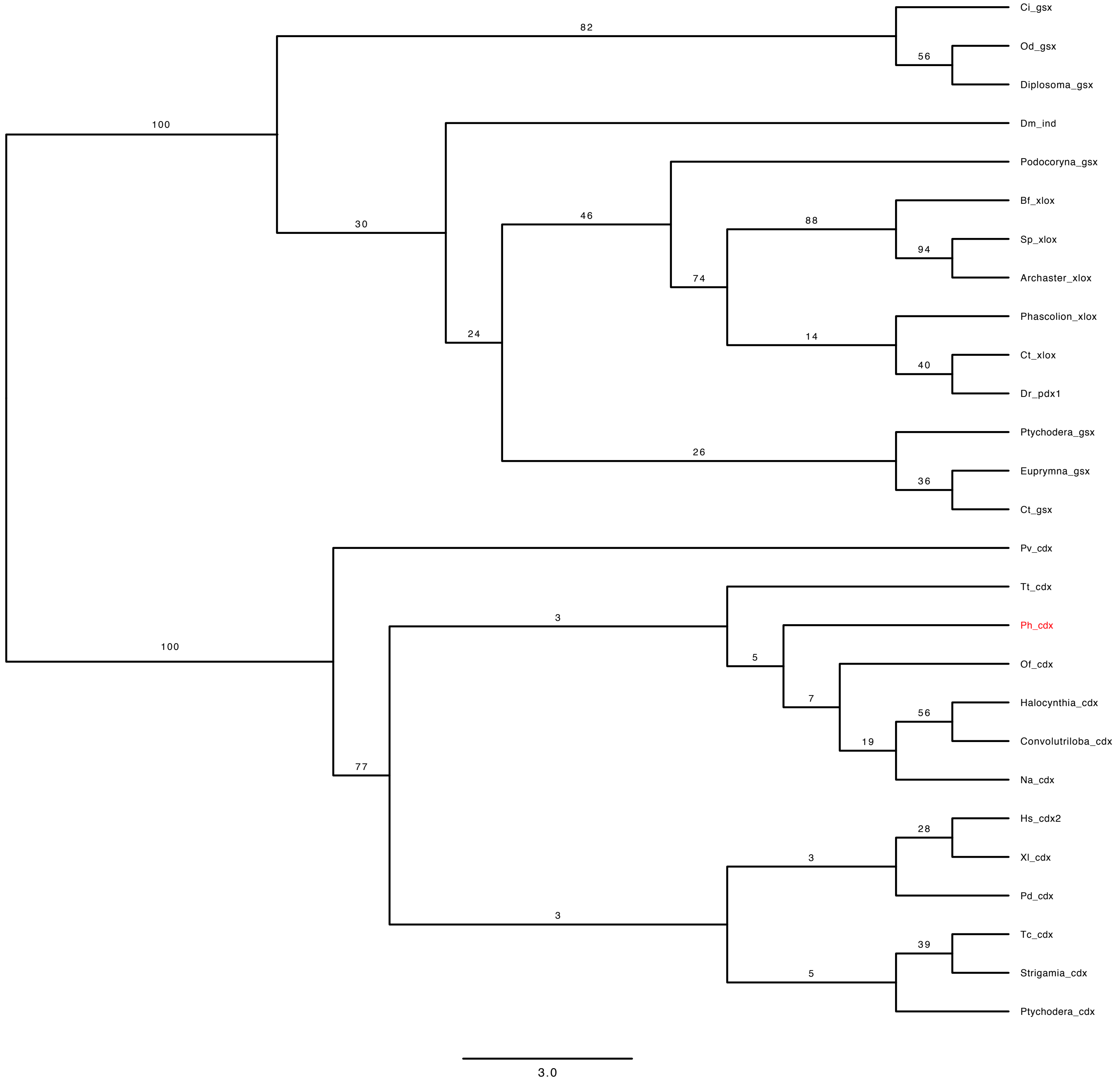

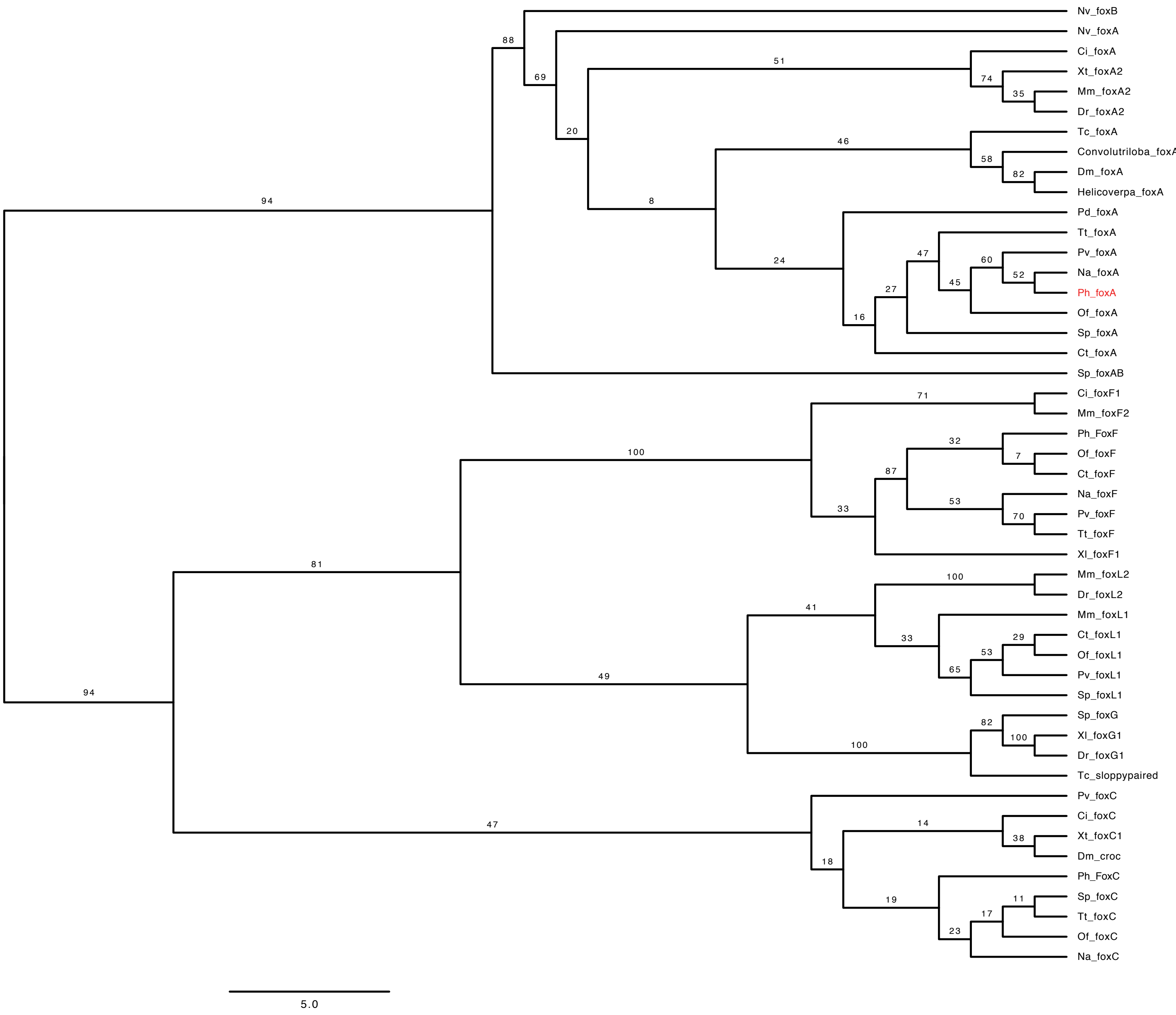

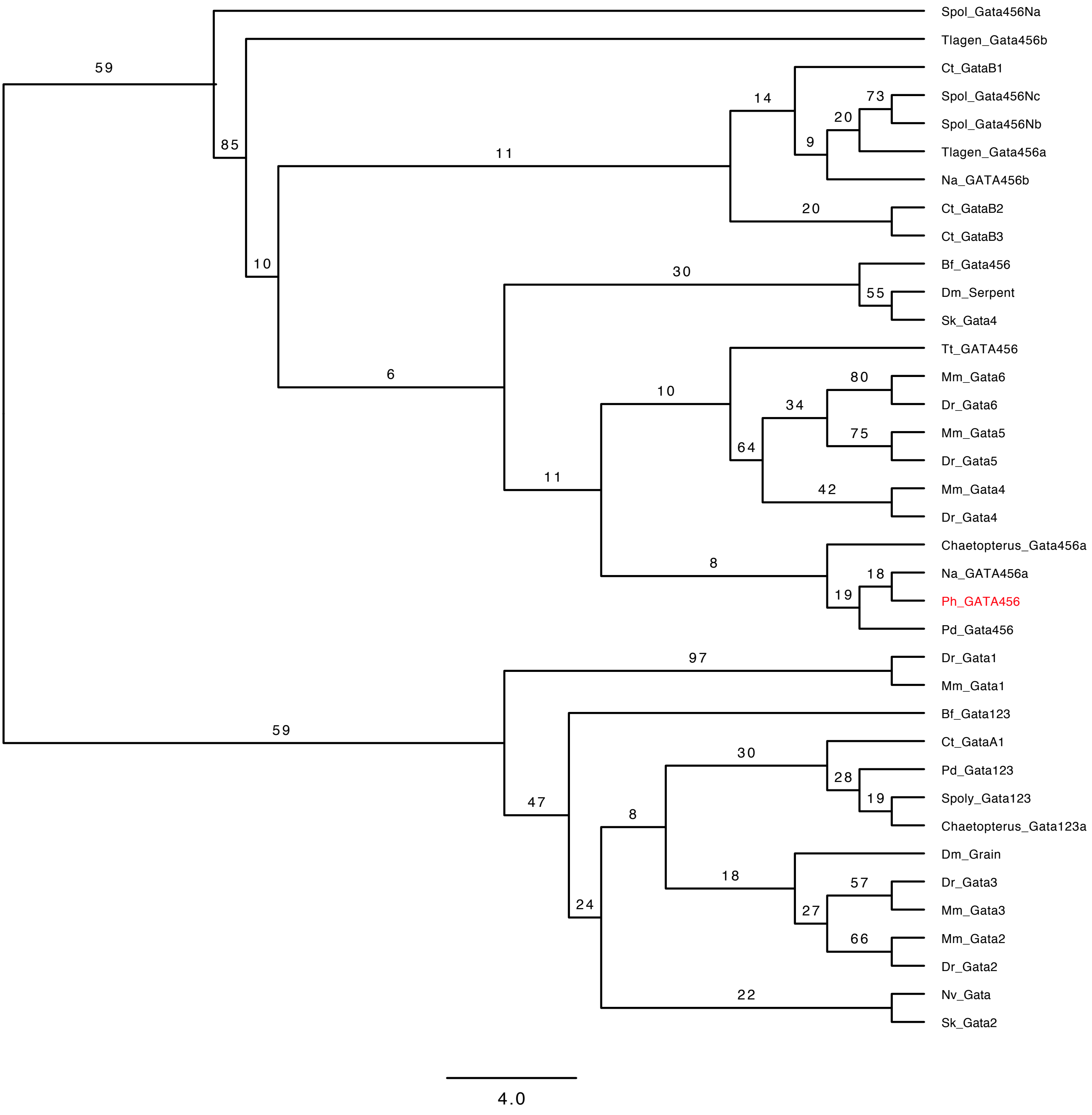

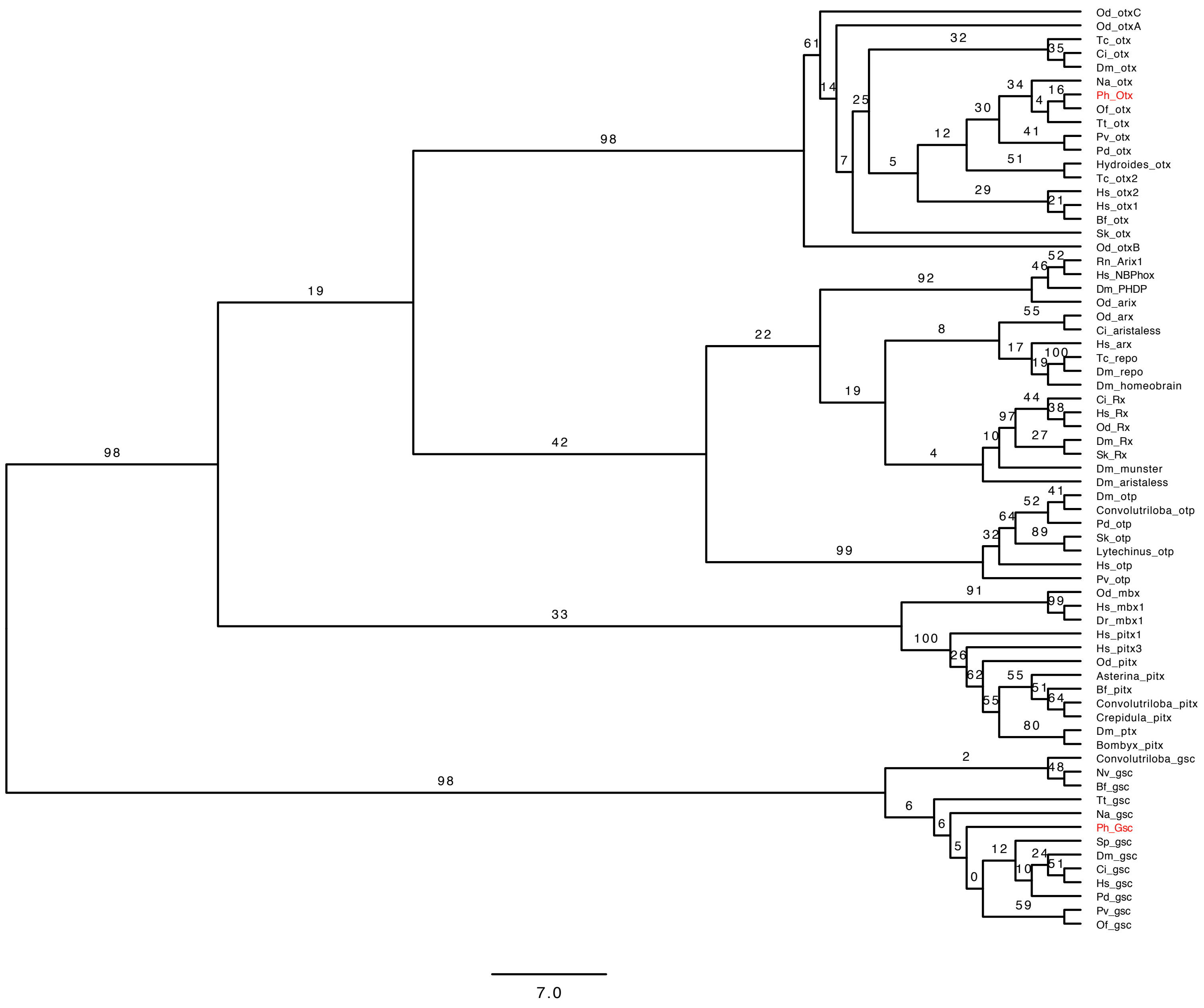

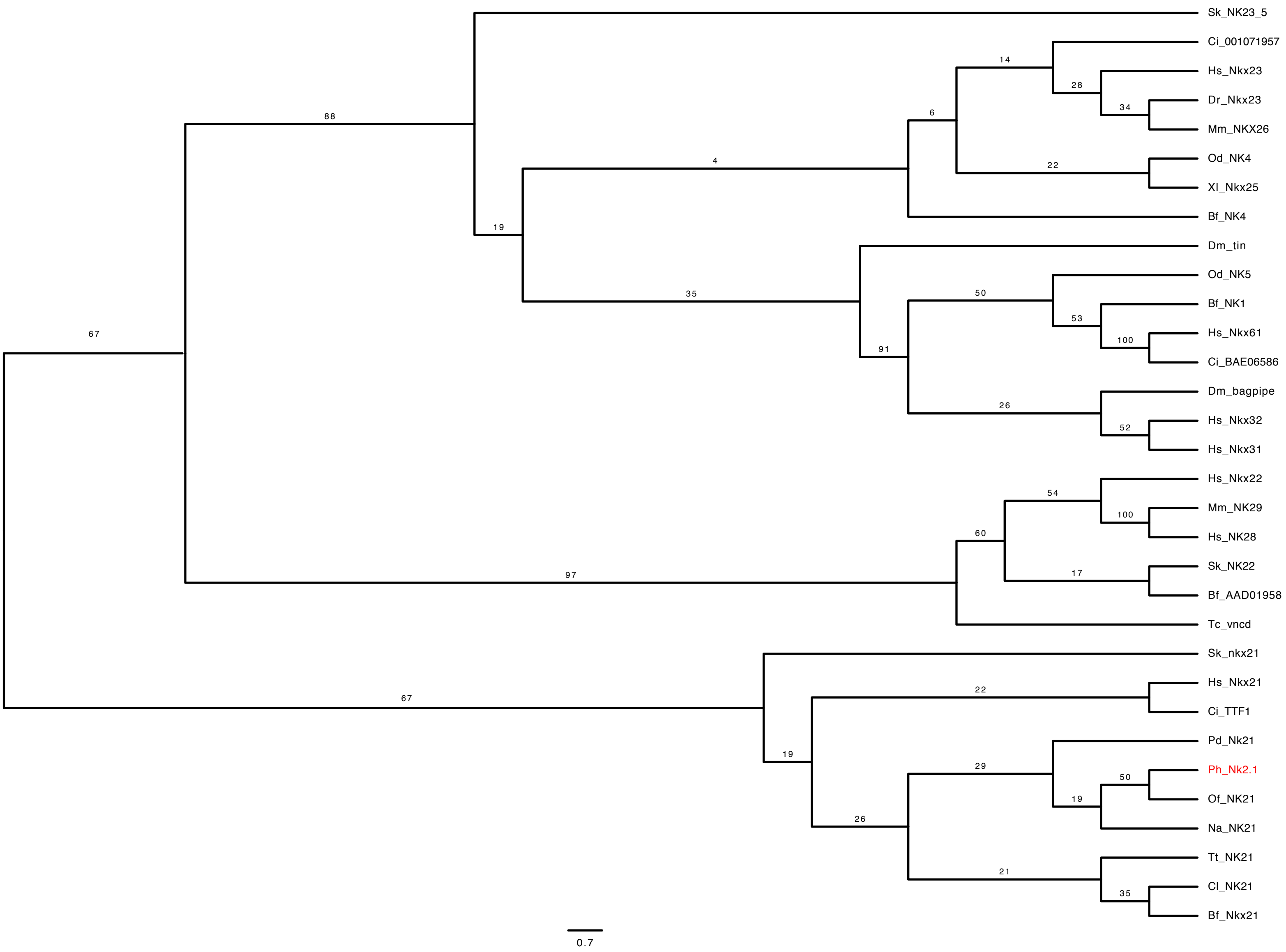

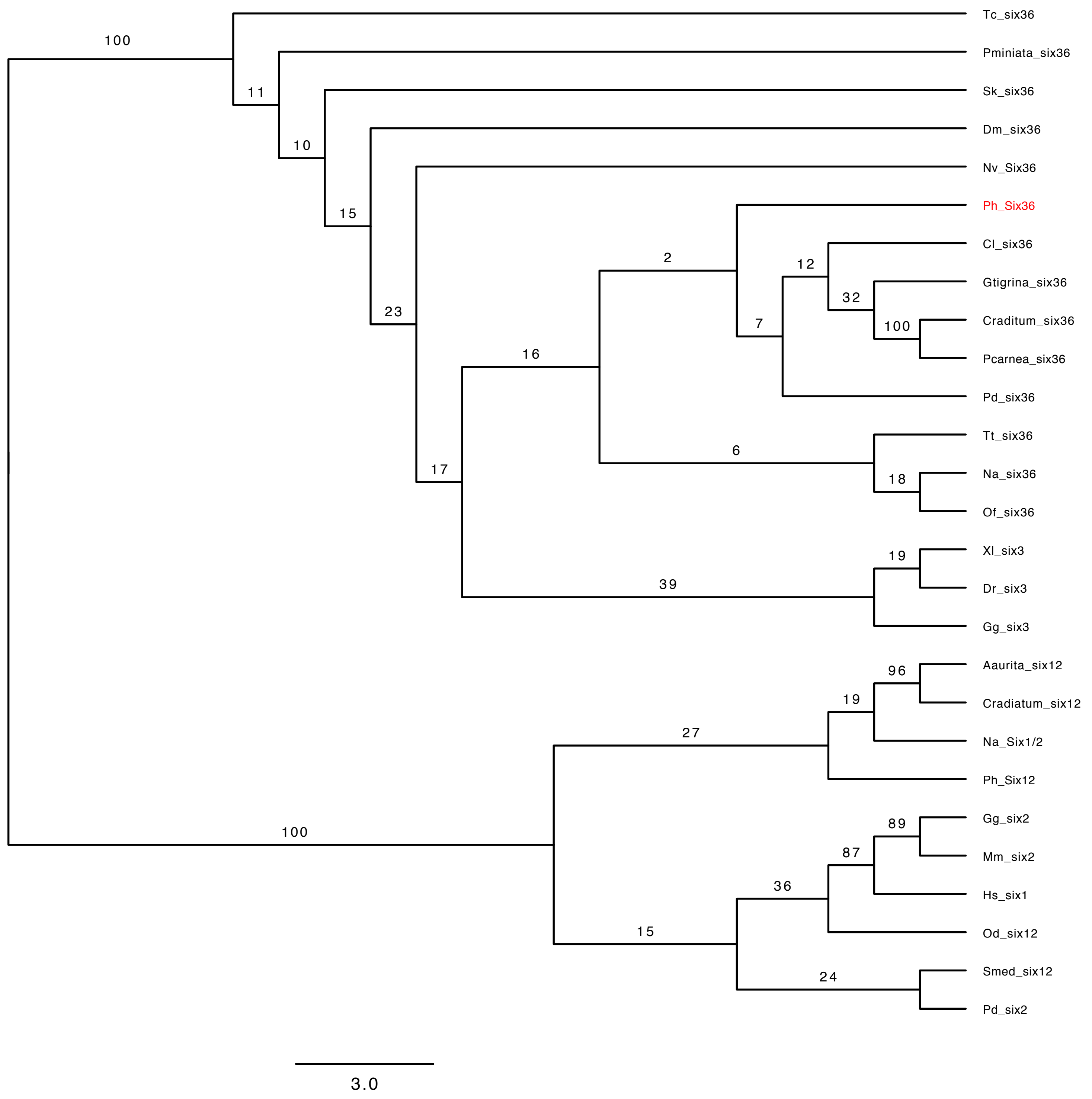

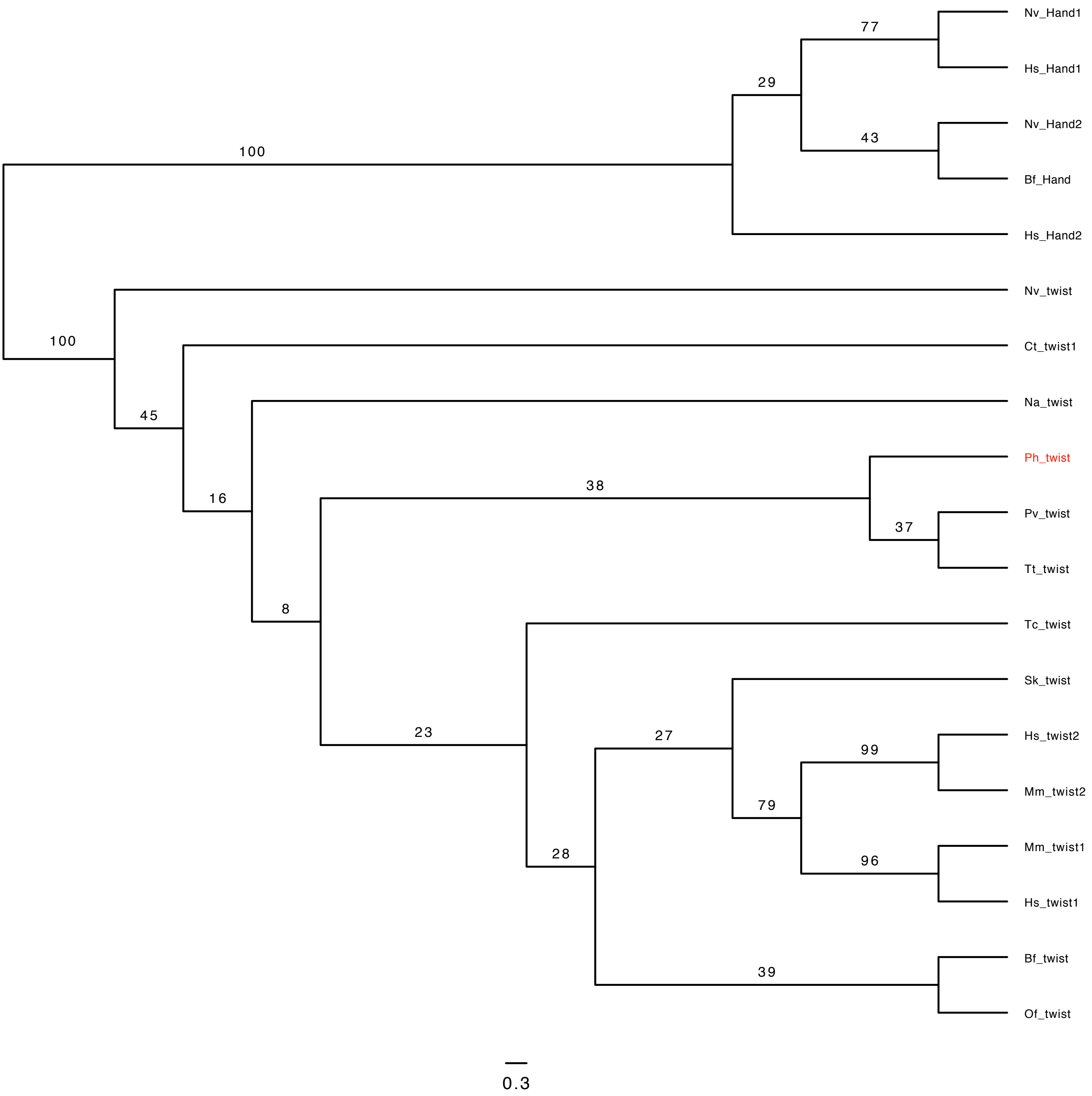
